# Immunological characterization of peritoneal exudate cells in liver cirrhosis patients

**DOI:** 10.1101/2025.02.19.638957

**Authors:** Shiori Kaji, Izumi Sasaki, Satoko Kunimoto, Naoko Wakaki-Nishiyama, Emi Tosuji, Ayano Kurara, Takashi Kato, Chisako Mutsukawa, Daisuke Okuzaki, Chizuyo Okamoto, Yukiko Yamano, Ryo Shimizu, Shuya Maeshima, Yoshiyuki Ida, Yasunobu Yamashita, Tadamichi Hashimoto, Sadahiro Iwabuchi, Shinichi Hashimoto, Shin-Ichiroh Saitoh, Shin-ichi Araki, Masayuki Kitano, Tsuneyasu Kaisho

## Abstract

**BACKGROUND AND AIMS:** Liver cirrhosis (LC) is the end stage of liver fibrosis caused by various chronic liver diseases. Patients with LC often develop ascites containing peritoneal exudate cells (PECs). However, those cells have not been fully immunologically characterized. In this study, we clarify immune cell profiles of PECs from patients with LC.

**APPROACH AND RESULTS:** PECs were collected from patients with LC or patients receiving continuous ambulatory peritoneal dialysis (CAPD) as a non-cirrhotic control and subjected to single-cell RNA sequencing (scRNA-seq), bulk RNA sequencing (RNA-seq) and flowcytometry analyses. Analysis of scRNA-seq revealed that dendritic cells (DCs) and macrophages were major populations in CAPD patient-derived PECs, while those cells were decreased and T cells were most abundant in LC patient-derived PECs. Notably, *FCGR3A*-expressing macrophages were dominant over DCs and *GATA6*-expressing macrophages in LC patient-derived PECs. Bulk RNA-seq analysis further clarified expression of a set of genes was up- or down-regulated along with LC severity. Especially, expression of T cell signature genes was featured by its increase at Child-Pugh class B, but decrease at Child-Pugh class C. Flowcytometry analysis showed increase of T cells, decrease of DCs and macrophages, and increased expression of CD16 and CD163 in CD1c^int^CD14^high^ cells corresponding to *FCGR3A*-expressing macrophages in LC patient-derived PECs.

**CONCLUSIONS:** In LC patient-derived PECs, myeloid and T cell populations and their gene expression profiles were fluctuated with severity. Our findings should contribute to further development of diagnosis or therapeutic maneuver for LC.

## INTRODUCTION

Liver cirrhosis (LC) is an end stage of liver fibrosis and causes more than one million deaths every year all over the world.^1^ As liver fibrosis progresses, patients often develop ascites, mainly due to portal hypertension. A variety of immune cells can be found in the ascites and they change their profiles, depending on the pathological state of the patients. For example, in patients with LC, CD8^+^ mucosal-associated invariant T (MAIT) cells and CD16^high^ CD163^high^ macrophages are increased in the ascites but not in peripheral bloods.^2^ In mice, peritoneal cavity macrophages migrate into the liver upon liver injury,^3^ and regulate the pathological conditions in the liver.^4,5^ These studies indicate peritoneal cavity cells should have critical roles in liver pathogenesis. However, it is unclear whether or how various types of immune cells fluctuate in the ascites of patients with LC.

In this study, we analyzed the peritoneal exudate cells (PECs) of patients with LC in comparison with those of patients undergoing continuous ambulatory peritoneal dialysis (CAPD). We then found increase in frequency of T cells and macrophage subsets and decrease in frequency of dendritic cells (DCs) in the ascites of LC patients. Furthermore, we have also revealed some parameters correlating with disease severity.

## MATERIALS AND METHODS

### Patients

Patients with LC and patients with end-stage renal disease undergoing CAPD at Wakayama Medical University were eligible for this study (Supplementary Table S1-3). Written informed consent was obtained from all patients, and the study protocol conforms to the ethical guidelines of the 2013 Declaration of Helsinki and the 2018 Declaration of Istanbul. All studies were performed in accordance with appropriate ethical regulations and were pre-approved by the Wakayama Medical University Research Ethics Committee (#3918).

### Preparation of peritoneal exudate cells

PECs from patients with LC were collected from ascites by puncture and drainage at the diagnosis or treatment. PECs from patients undergoing CAPD were collected from 1.5-2 L of CAPD effluent according to the standard CAPD prescription. Ascites of patients with LC or CAPD effluent were centrifuged at 4℃ 2000 rpm 5 min. Cell pellets were washed three times with phosphate buffered saline (PBS) and resuspended in PBS supplemented with 1% fetal calf serum (FCS), 5 mM ethylenediaminetetraacetic acid (EDTA) and 0.01% sodium azide (NaN_3_). Turk’s solution was used to analyze cell viability to ensure single-cell suspension with more than 90% viability.

### Single-cell RNA sequencing analysis

Single-cell RNA sequencing (scRNA-seq) analysis was performed using Chromium Controller (10x Genomics, Pleasanton, CA). Libraries were sequenced on NovaSeq 6000 (Illumina, San Diego, CA) in a 101+101 base paired-end mode. Illumina RTA v3.4.4 software was used for base calling. Generated reads were input into the cellranger pipeline (cellranger-7.1.0, 10x Genomics) for UMI counting for each gene-cell (barcode) combination. The cluster analysis based on Uniform Manifold Approximation and Projection (UMAP) plot and differential expression analysis were performed by Loupe Browser (v7.0.1, 10x Genomics).

### Bulk RNA sequencing analysis

Total RNA was extracted with RNeasy micro kit (QIAGEN, Hilden, Germany). Full-length cDNA was prepared using a SMART-Seq HT Kit (Takara Bio, Japan). According to the SMART-Seq kit instructions, an Illumina library was then prepared using a NexteraXT DNA Library Preparation Kit (Illumina). Sequencing was performed on NovaSeq 6000 platform in a 101-base single-end mode. Illumina RTA v3.4.4 software was used for base calling. Generated reads were mapped to the human (hg19) reference genome using TopHat v2.1.1 in combination with Bowtie2 ver. 2.2.8 and SAMtools ver. 0.1.18. Fragments per kilobase of exon per million mapped fragments (FPKMs) were calculated using Cuffdiff 2.2.1 with parameter -max-bundle-frags 50000000. The DAVID Bioinformatics database and Kyoto Encyclopedia of Genes and Genomes (KEGG) annotations were used for enrichment analysis. The web tools ClustVis and VolcaNoseR^6^ were used to draw heatmaps and volcano plots, respectively.

### Flowcytometry analysis

PECs were collected in Eppendorf tubes from patients undergoing CAPD or patients with LC, and subjected to flowcytometry analysis. PECs were stained with the antibodies listed in Supplementary Table S4. Dead cells detected as positive for AmCyan channels were excluded using the LIVE/DEAD Fixable Dead Cell Aqua Stain Kit (cat; L34966, Invitrogen, Waltham, MA). Stained cells were analyzed with FACS Verse and data were processed with FlowJo Version 10.8.1 software (TreeStar, Ashland, OR).

### Magnetic beads cell isolation

CD1c^high^ CD14^int^, CD1c^int^ CD14^high^ or CD1c^low-int^ CD14^low^ cells were isolated from PECs by using magnetic-activated cell sorting (MACS). PECs were stained with biotinylated anti-CD1c antibody for 20 min on ice. After washing, stained PECs were applied to streptavidin microbeads (Miltenyi Biotec, Bergisch Gladbach, Germany) and then magnetically sorted with LS columns as per the manufacturer’s instructions. Streptavidin microbeads-conjugated cells were used as CD1c^high^ CD14^int^ cells. The flow-through cells including CD1c^int^ CD14^high^ and CD1c^low-int^ CD14^low^ cells were stained with PE-conjugated anti-CD14 antibody for 20 min on ice. After washing, cells were applied to PE microbeads (Miltenyvi Biotec) and then magnetically sorted with LS columns as per the manufacturer’s instructions. PE microbeads-conjugated cells were used as CD1c^int^ CD14^high^ cells and the flow-through cells were used as CD1c^low-int^ CD14^low^ cells. Purity of CD1c^high^ CD14^int^, CD1c^int^ CD14^high^ or CD1c^low-int^ CD14^low^ cells were verified by flowcytometry analysis.

### Quantitative real-time PCR

Using PrimeScript RT Reagent Kit (TaKaRa, Kusatsu, Japan), 0.5 μg of total RNA was reverse transcribed into complementary DNA. Relative expression of RNA transcripts was quantified using gene-specific primers, TB Green Premix Ex Taq II (TaKaRa) and the StepOnePlus Real-Time PCR System (Applied Biosystems, Waltham, MA). TaqMan probes (TaqMan Gene Expression Assay, Applied Biosystems) were used for 18S rRNA (internal control). Expression of all genes was normalized to that of 18S rRNA and is represented as the ratio to the indicated reference samples. Gene-specific primers used in the study were obtained from FASMAC, and all primers were validated for linear amplification. The primer sequences were designed and synthesized by FASMAC as follows: *GATA6*, forward primer: 5’- GCCACTACCTGTGCAACGCCT-3’, reverse primer: 5’-CAATCCAAGCCGCCGTGATGAA- 3’.

### Statistical analysis

Statistical significance was determined by an unpaired two-tailed Student’s t test. *p* values are indicated by \**p* < 0.05, \*\**p* < 0.01, \*\*\**p* < 0.001, \*\*\*\**p* < 0.0001. *p* < 0.05 was considered statistically significant.

### Data availability statement

Bulk RNA sequencing (bulk RNA-seq) and scRNA-seq data generated in this study have been deposited at GEO under accession number GSE287363 and GSE287364, respectively.

## RESULTS

### Single-cell RNA-seq analysis of LC and CAPD patient-derived PECs

We first collected PECs from a patient with LC at Child-Pugh class B (CP-B) and from a patient undergoing CAPD on the same day and performed scRNA-seq analysis (Fig. 1A). According to several signature genes, five cell clusters including T cells, B cells, *GATA6*-expressing macrophages, DCs and *FCGR3A*-expressing macrophages were defined (Fig. 1B,C). In CAPD patient-derived PECs, the frequencies of T cells, B cells, *GATA6*-expressing macrophages, DCs and *FCGR3A*-expressing macrophages were 18%, 1.4%, 9.8%, 30% and 40%, respectively. DCs and macrophages occupied around 80% of total PECs. In LC patient-derived PECs, T cells were significantly increased to 73% and became the most abundant population. Meanwhile, *GATA6*-expressing macrophages, DCs and *FCGR3A*-expressing macrophages were decreased to 1.6%, 2.1% and 16%, respectively. Due to the most prominent decrease of DCs, *FCGR3A*-expressing macrophages were dominant over *GATA6*-expressing macrophages and DCs in LC patient-derived PECs.

**FIG. 1.**
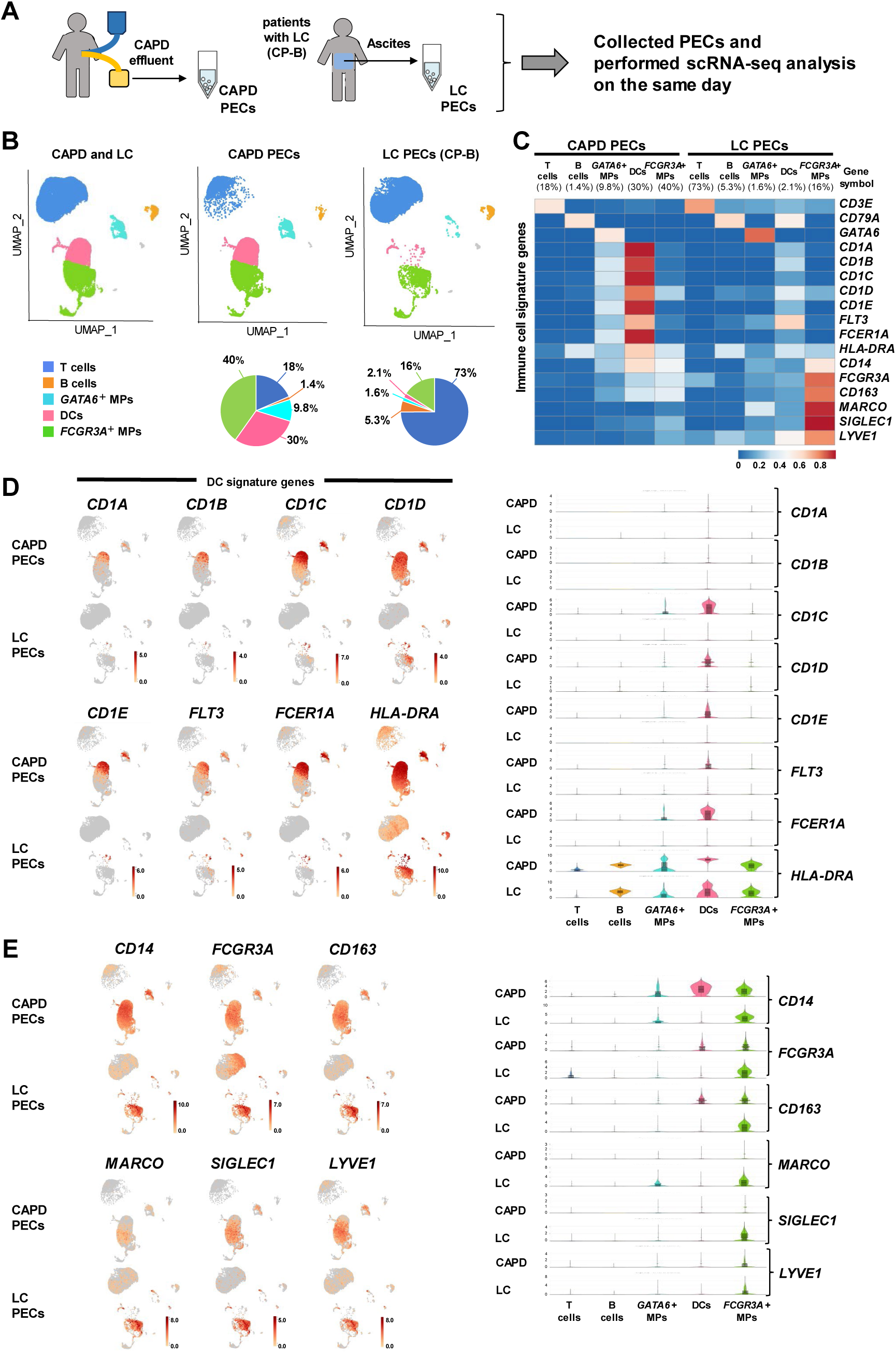
Single-cell RNA sequencing analysis revealed major cell clusters and gene expression profiles of CAPD and LC patient-derived PECs. (A) Schematic workflow of scRNA-seq analysis of human PECs. (B) UMAP shows five distinct clusters of CAPD PECs (CAPD patient-derived PECs) and LC PECs (LC patient-derived PECs). Clusters include T cells, B cells, *GATA6*-expressing macrophages (*GATA6*^+^ MPs), DCs and *FCGR3A*-expressing macrophages (*FCGR3A*^+^ MPs). (C) Heatmap shows five clusters of CAPD and LC PECs and the expression of immune cell signature genes. (D) Feature plots and violin plots of DC signature gene expression in CAPD and LC PECs. (E) Feature plots or violin plots of expression of *CD14*, *FCGR3A*, *CD163*, *MARCO*, *SIGLEC1* and *LYVE1* in CAPD and LC PECs.

We then investigated expression of DC signature genes. According to the heatmap, feature plot and violin plot analyses, expression of DC signature genes, including CD1 family members (*CD1A, CD1B, CD1C, CD1D, CD1E*), *FLT3*, *FCER1A* and HLA family gene (*HLA-DRA*), were much lower in DCs of LC patient-derived PECs than in DCs of CAPD patient-derived PECs (Fig. 1C,D).

*FCGR3A*-expressing macrophages are featured by expression of *CD163*, *MARCO*, *SIGLEC1* and *LYVE1*, and this type of macrophage has been reported to be involved in tissue repair.^7^ *CD163*, *MARCO*, *SIGLEC1* and *LYVE1* were expressed in *FCGR3A*-expressing macrophages in CAPD patient-derived PECs (Fig. 1C,E). Expression levels of these genes was increased in LC patient-derived PECs.

In mice, macrophages expressing a transcription factor, GATA6, occupy around 90% of resident peritoneal macrophages.^8–10^ GATA6 is essential for generation of resident peritoneal macrophages and its target genes include *Serpinb2*, *Nt5e*, *Itga6*, *Thbs1*, *Tgfb2*, *Ltbp1*^10^ and *CD9*.^9^ *GATA6*-expressing macrophages also exist in humans, although they represent less than 10% and are not a major population.^11^ *CD9*, *SERPINB2, NT5E, ITGA6, THBS1, TGFB2* and *LTBP1* were highly expressed in *GATA6*-expressing macrophages in CAPD patient-derived PECs (Fig. 2A-C). Expression levels of all of these genes except *CD9* and *NT5E* were decreased, while *CD9* and *NT5E* expression was increased in LC patient-derived PECs. *PDGFRB*, *KRT5*, *ACTA2*, *COL1A1* and *UPK3B* were expressed in peritoneal mesothelial cells and classified as peritoneal mesothelial cell signature genes.^12^ All of these genes were expressed selectively in *GATA6*-expressing macrophages in CAPD patient-derived PECs (Fig. 2D-F). Furthermore, among them, *PDGFRB*, *KRT5*, *ACTA2*, *COL1A1* and *UPK3B* were expressed at higher levels in LC than in CAPD patient-derived PECs.

**FIG. 2.**
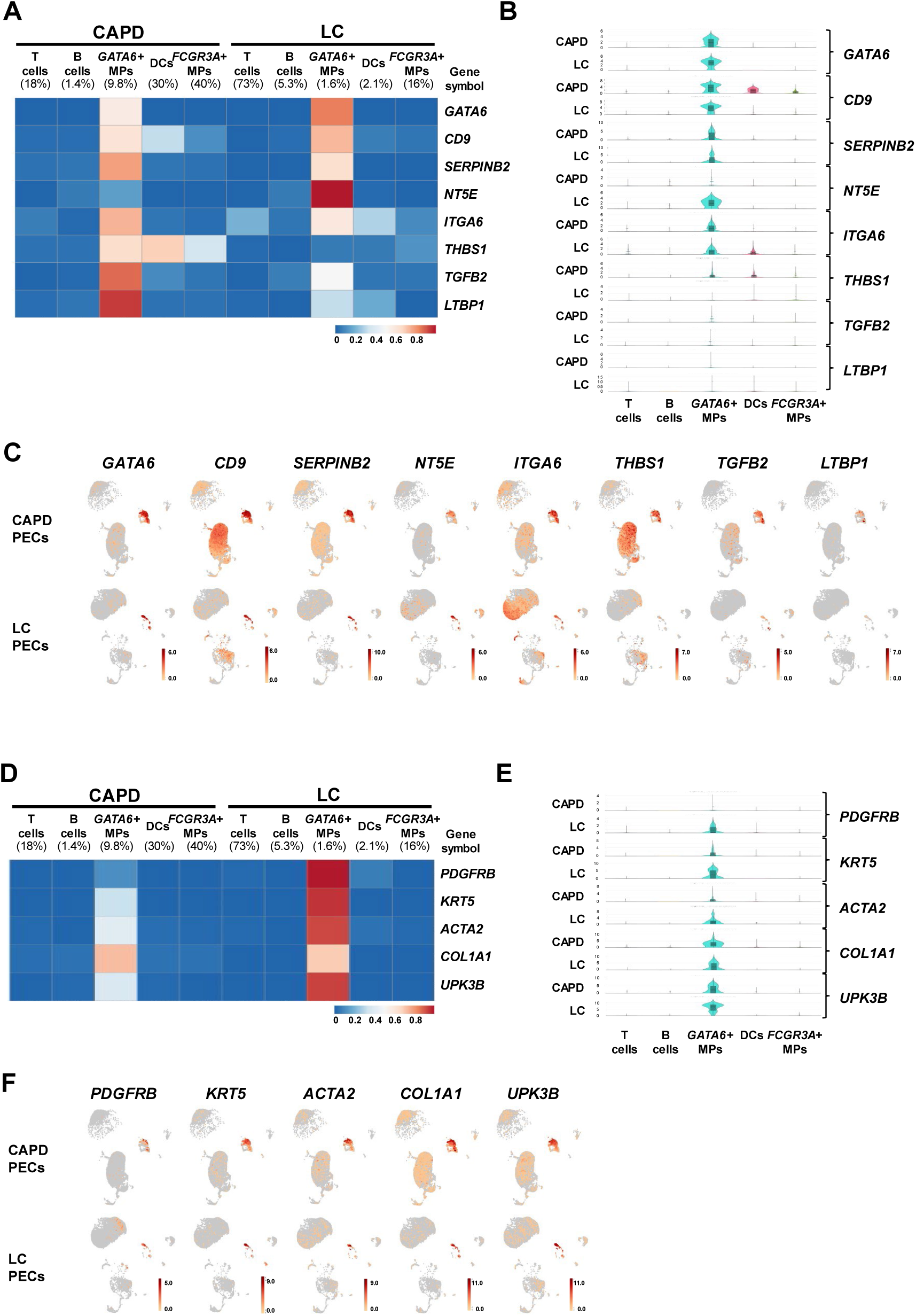
Single-cell RNA sequencing analysis revealed gene expression profiles of *GATA6*-expressing macrophages in CAPD and LC patient-derived PECs. (A-C) Gene expression of *GATA6*, *CD9*, *SERPINB2*, *NT5E*, *ITGA6*, *THBS1*, *TGFB2* and *LTBP1* is shown in heatmap (A), violin plots (B) and feature plots (C). (D-F) Gene expression of *PDGFRB*, *KRT5*, *ACTA2*, *COL1A1* and *UPK3B* is shown in heatmap (D), violin plots (E) and feature plots (F).

We then analyzed T cell populations. In LC patient-derived PECs, T cell percentages among PECs were increased and *CD8A*^+^ T cells dominated over *CD4*^+^ T cells, but the ratio of *CD4*^+^ to *CD8A*^+^ T cells were not so different from that of CAPD patient-derived PECs (Fig. 3A-C). In both LC and CAPD patient-derived PECs, *CD8A*^+^ T cells included a significant fraction of cells expressing *CD8A*, *TRAV1-2* (encoding TCR Vα7.2) and *KLRB1* (encoding CD161), which represent MAIT cells.

**FIG. 3.**
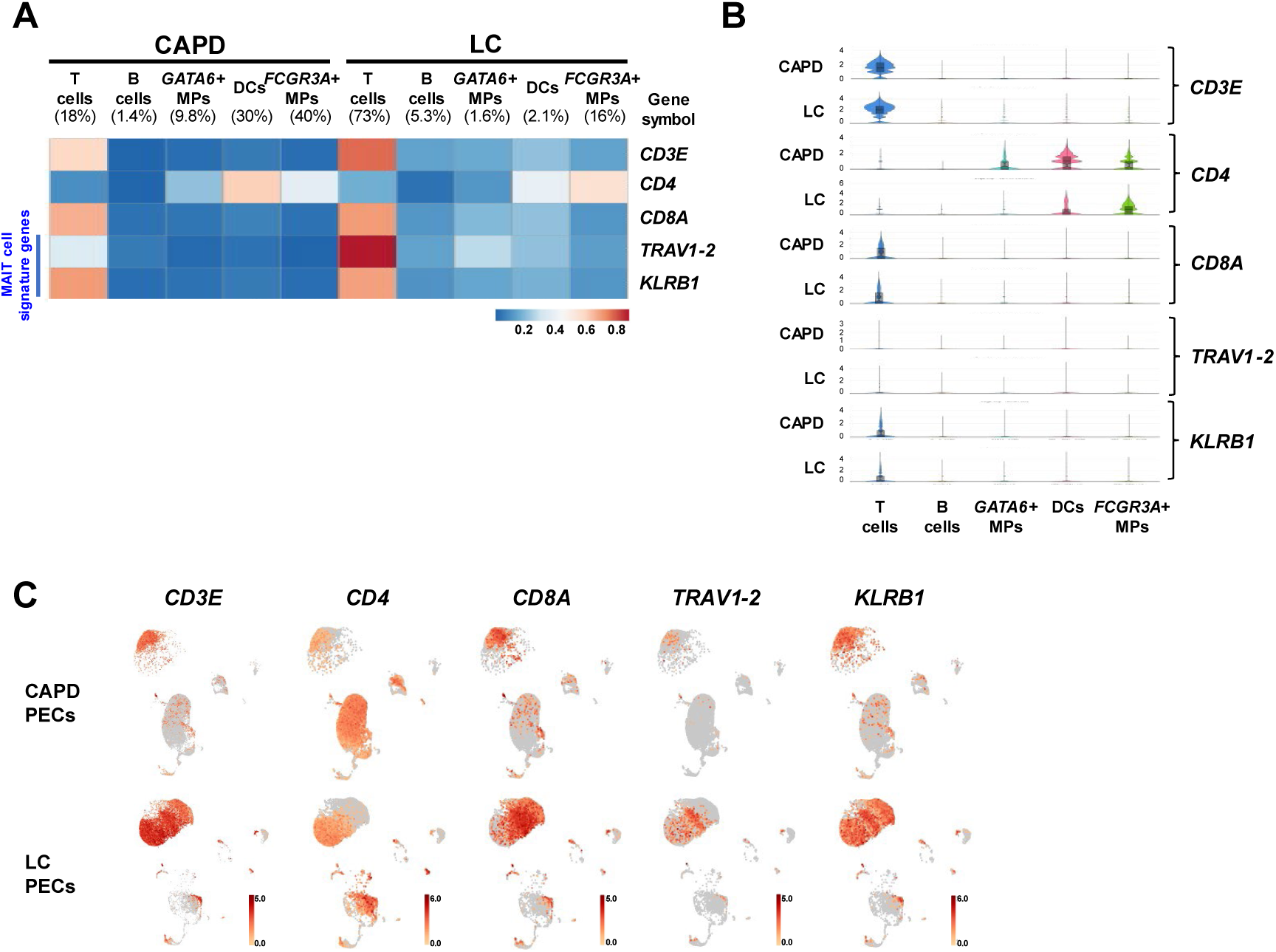
Single-cell RNA sequencing analysis revealed gene expression profiles of T cells in CAPD and LC patient-derived PECs. (A-C) Gene expression of *CD3E*, *CD4*, *CD8A*, *TRAV1-2* and *KLRB1* is shown in heatmap (A), violin plots (B) and feature plots (C).

### Bulk RNA-seq analysis revealed down- or up-regulated genes in LC patient-derived PECs

We also performed bulk RNA-seq analysis in PECs from patients with LC and patients undergoing CAPD (Fig. 4A). In comparison to PECs from patients undergoing CAPD, expression of 1107 and 906 genes were more than 2-fold down- and up-regulated, respectively, with significance in PECs from patients with LC (Fig. 4B). The top five downregulated genes in PECs from patients with LC included DC signature genes, such as *CD1C*, *CD1E*, *CD1B* and *FCER1A* (Fig. 4C).

**FIG. 4.**
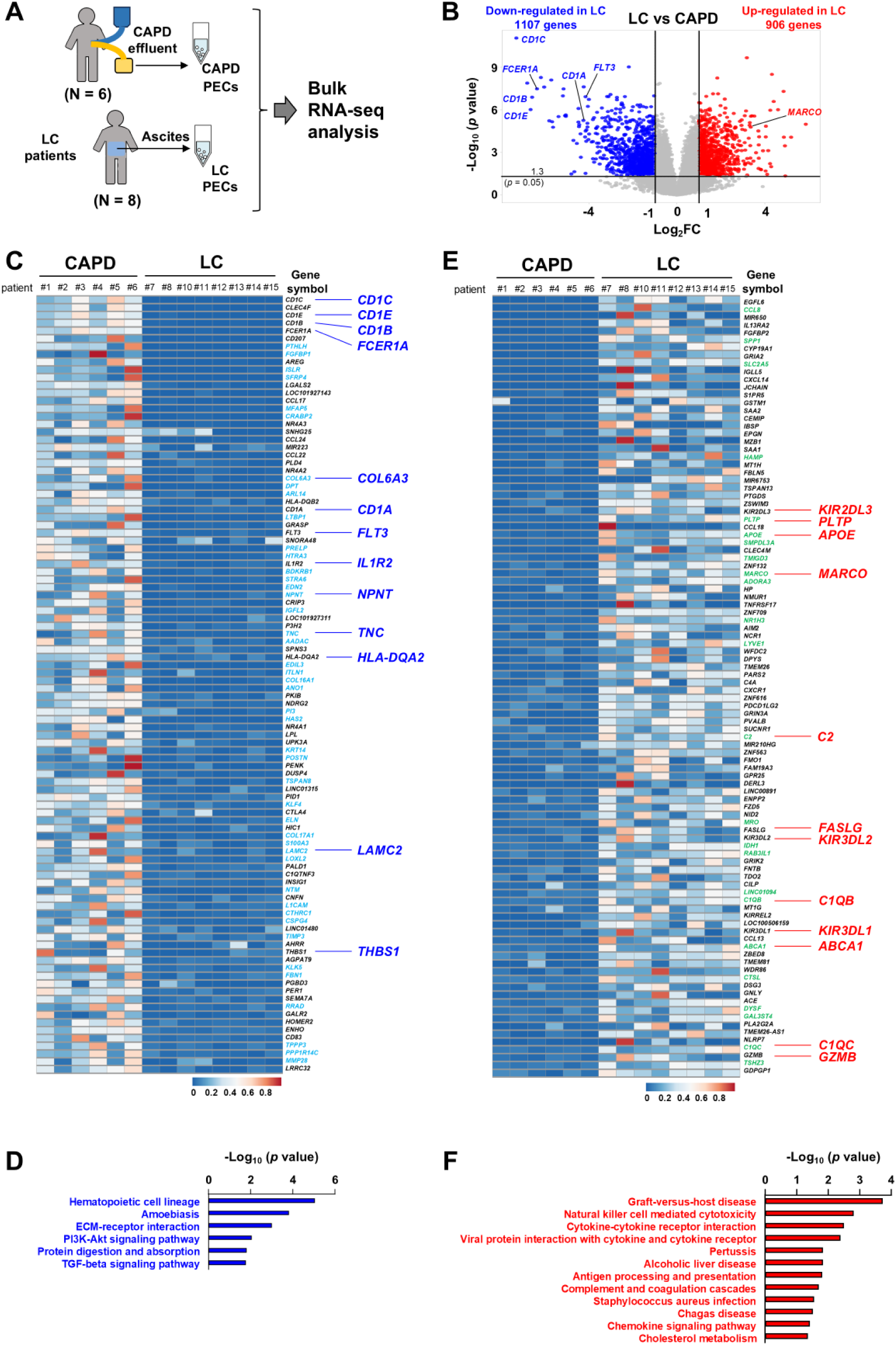
Bulk RNA sequencing analysis revealed that DCs and *GATA6*-expressing macrophages mainly decreased in LC patient-derived PECs. (A) Schematic workflow of bulk RNA-seq analysis of human PECs. (B) Volcano plots of gene expression in PECs show a fold change (FC) and *p* value for the comparison of LC PECs versus CAPD PECs. Up- or down-regulated genes (FC ≥ 2, *p* < 0.05) are depicted in red or blue, respectively. (C) Heatmap of the top 100 downregulated genes in LC PECs. (D) KEGG pathway analysis of downregulated genes in (C). (E) Heatmap of the top 100 upregulated genes in LC PECs. (F) KEGG pathway analysis of upregulated genes in (E).

KEGG pathway analysis of the top 100 downregulated genes in PECs from patients with LC showed that ‘Hematopoietic cell lineage’ was the most enriched pathway (Fig. 4D). The pathway contained CD1 family genes (*CD1A*, *CD1B*, *CD1C* and *CD1E*) and *FLT3*, *IL1R2* and *HLA-DQR2*, all of which are dominantly expressed in DCs (Supplementary Figure 1A,B). These CD1 family genes (*CD1A*, *CD1B*, *CD1C* and *CD1E*), *IL1R2* and *LAMC2* were also contained in the second enriched pathway, ‘Amoebiasis’. The third enriched pathway was ‘ECM-receptor interaction’ pathway, which contained *TNC*, *NPNT*, *COL6A3* and *THBS1*. Among the four genes, *TNC*, *NPNT* and *COL6A3* were dominantly expressed in *GATA6*-expressing macrophages and *THBS1* was expressed in DCs and *GATA6*-expressing macrophages. According to the scRNA-seq analysis, we categorized the top 100 downregulated genes in PECs from patients with LC. 82 genes were expressed in DCs or *GATA6*-expressing macrophages. Among them, 37 genes were expressed in both DCs and *GATA6*-expressing macrophages. 45 genes were dominantly expressed in *GATA6*-expressing macrophages, while no genes were expressed dominantly in DCs (shown in light blue in Fig. 4C). Thus, the gene expression analysis indicates that DCs and *GATA6*-expressing macrophages mainly decreased in PECs from patients with LC.

KEGG pathway analysis of 100 upregulated genes in PECs from patients with LC showed ‘Graft-versus-host disease’ was the most enriched pathway (Fig.4E, F). This pathway contained *GZMB*, *KIR3DL1*, *FASLG*, *KIR3DL2* and *KIR2DL3*, which are highly expressed in T cells (Supplementary Figure 1C,D). Among 100 upregulated genes, 24 genes were dominantly expressed in *FCGR3A*-expressing macrophages (shown in green in Fig. 4E). Those genes include complement genes such as *C1QB*, *C1QC* and *C2*, which belong to the ‘Alcoholic liver disease’ pathway. They also include *ABCA1*, *APOE* and *PLTP*, which belong to the ‘Cholesterol metabolism’ pathway. *C1QB*, *C1QC*, *C2*, *ABCA1*, *APOE* and *PLTP* were highly expressed in *FCGR3A*-expressing macrophages in PECs from patients with LC (Supplementary Figure 1C,D). These results indicate that T cells and *FCGR3A*-expressing macrophages were increased in PECs from patients with LC.

### Comprehensive gene expression analysis in correlation with LC severity

We then performed comprehensive gene expression analysis in correlation with LC severity (Fig. 5A, left). Severity of LC was assessed by Child-Pugh (CP) classification, which are widely used.^13^ Among eight patients with LC, four patients, including a patient subjected to scRNA-seq analysis, were CP-B and the other 4 patients were Child-Pugh class C (CP-C).

**FIG. 5.**
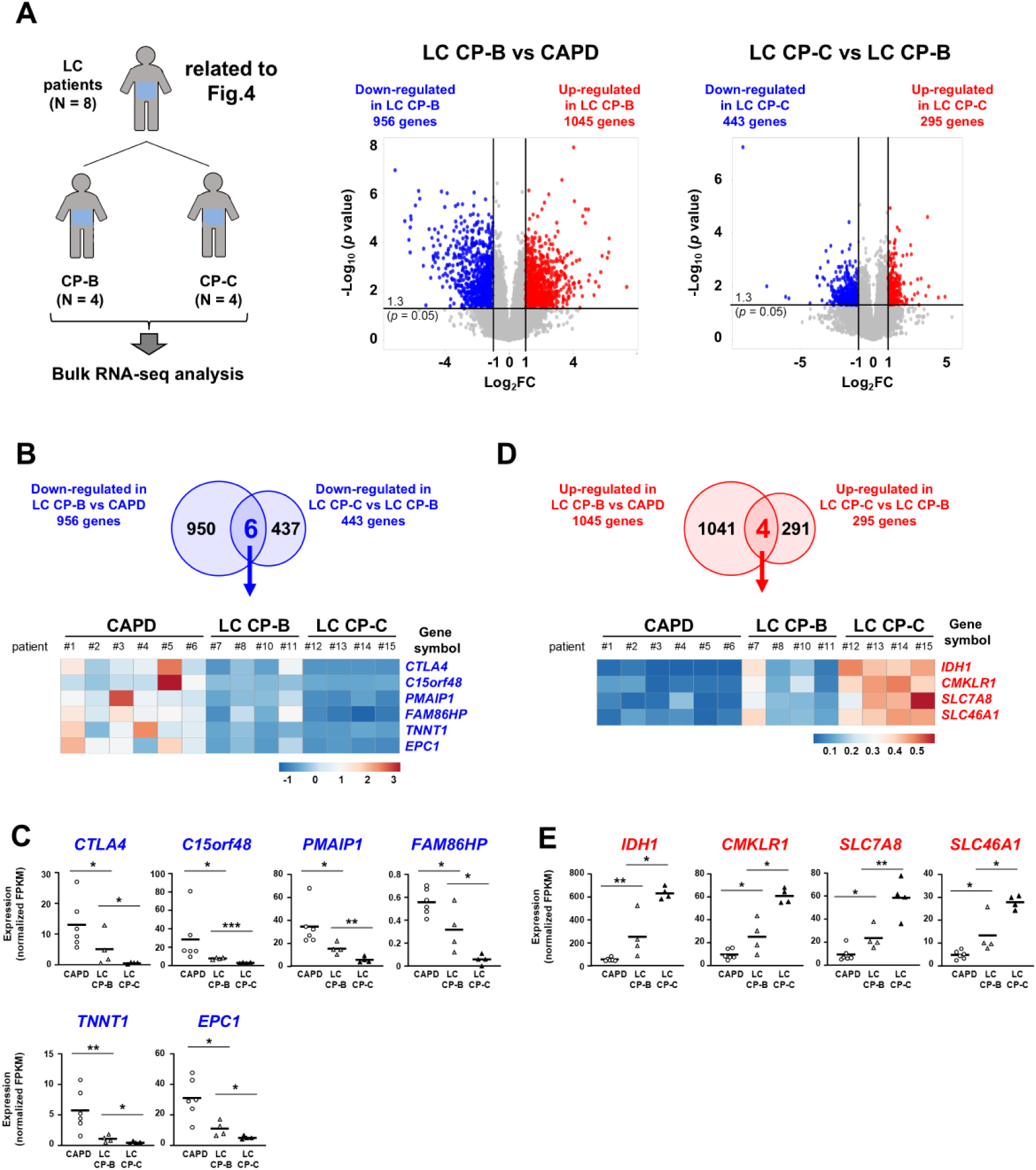
Bulk RNA sequencing analysis revealed up- or down-regulated genes in correlation with LC severity. (A) Volcano plots of gene expression show a FC and *p* value for the comparisons of LC CP-B PECs versus CAPD PECs (middle) or LC CP-C PECs versus LC CP-B PECs (right). Two-fold up- or down-regulated genes (FC ≥ 2, *p* < 0.05) with significance are depicted in red or blue, respectively. (B) Venn diagram depicting the numbers of 2-fold downregulated genes with significance in LC CP-B PECs (A, middle) or LC CP-C PECs (A, right), then expression of the overlapped six genes showed in heatmap. (C) Expression of the six genes (B). Statistical significance was determined by an unpaired two-tailed Student’s t test. *p* values are indicated by \**p* < 0.05, \*\**p* < 0.01. *p* < 0.05 was considered statistically significant. (D) Venn diagram depicting the numbers of 2-fold upregulated genes with significance in LC CP-B PECs (A, middle) or LC CP-C PECs (A, right), then expression of the overlapped four genes showed in heatmap. (E) Expression of the four genes (D). Statistical significance was determined by an unpaired two-tailed Student’s t test. *p* values are indicated by \**p* < 0.05, \*\**p* < 0.01. *p* < 0.05 was considered statistically significant.

We first searched the genes, expression of which decreased with disease severity. Expression of 956 genes were more than 2-fold downregulated in PECs from patients with LC CP-B, compared with PECs from patients undergoing CAPD (Fig.5A, middle). Expression of 443 genes were more than 2-fold downregulated in PECs from LC CP-C patients, compared with PECs from patients with LC CP-B (Fig.5A, right). Comparison of these genes revealed that expression of six genes (*CTLA4, C15orf48, PMAIP1, FAM86HP, TNNT1,* and *EPC1*) was decreased as LC gets worse (Fig. 5B,C). Among these genes, *CTLA4, C15orf48, PMAIP1* and *EPC1* were highly expressed in DCs and *GATA6*-expressing macrophages, while *TNNT1* was highly expressed in DCs and *FCGR3A*-expressing macrophages (Supplementary Figure 2A,B). We then searched the genes, the expression of which increased with disease severity.

Expression of 1045 genes were more than 2-fold upregulated in PECs from patients with LC CP-B, compared with PECs from patients undergoing CAPD (Fig.5A, middle). Expression of 295 genes were more than 2-fold upregulated in PECs from patients with LC CP-C, compared with PECs from patients with LC CP-B (Fig.5A, right). Comparison of these genes clarified that expression of four genes (*IDH1, CMKLR1, SLC7A8* and *SLC46A1*) was increased as LC becomes more severe (Fig.5D, E). All four of these genes were highly expressed in *GATA6*- and *FCGR3A*-expressing macrophages (Supplementary Figure 2C,D).

### T cell signature genes are downregulated in PECs from patients with LC CP-C

We have also searched genes, expression of which is upregulated at CP-B and then downregulated at CP-C. Expression of 126 genes were more than 2-fold upregulated in PECs from patients with LC CP-B, compared with PECs from patients with LC CP-C and PECs from patients undergoing CAPD (Fig.6 A,B). KEGG pathway analysis of these 126 genes showed ‘Natural killer cell mediated cytotoxicity’ pathway was enriched with significance (Fig.6B, C). Expression of *GZMB*, *IFNG*, *SH2D1A* and *LCK* in the pathway was elevated in PECs from patients with LC CP-B. According to scRNA-seq analysis, these four genes were highly expressed in T cells (Supplementary Figure 3A,B). These results suggest that T cells increased in PECs from patients with LC CP-B, and then decreased in PECs from patients with LC CP-C.

**FIG. 6.**
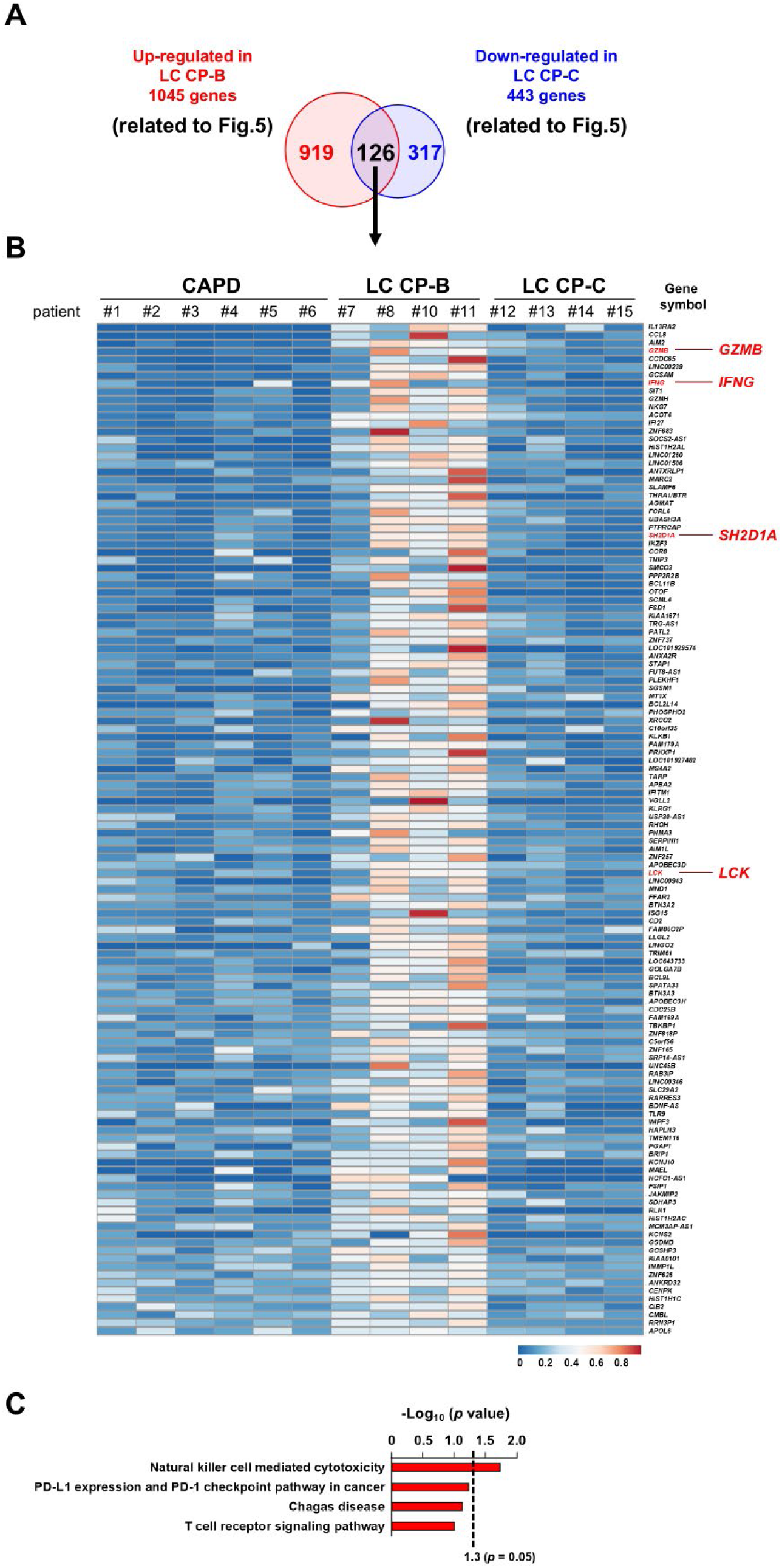
Bulk RNA sequencing analysis revealed that T cell signature genes were upregulated in LC CP-B but not LC CP-C. (A) Venn diagram depicting the numbers of upregulated genes in LC CP-B (Fig.5A, middle) or those of downregulated genes in LC CP-C (Fig.5A, right). (B) Heatmap of overlapped genes in (A). (C) KEGG pathway analysis of overlapped genes in (A).

### Decrease of CD1c^high^ CD14^int^ DCs and increase of CD8α^+^ MAIT cells in LC patient-derived PECs

We analyzed surface expression of several lineage markers in PECs. CD45^+^ cells of PECs are shown as the CD1c versus CD14 profile (Fig.7A). This profile showed three major clusters, CD1c^high^ CD14^int^ cells, CD1c^int^ CD14^high^ cells, and CD1c^low-int^ CD14^low^ cells. According to high expression of CD1c, CD1c^high^ CD14^int^ cells can be regarded as DCs. CD1c^int^ CD14^high^ cells can be regarded as *FCGR3A*-expressing macrophages, because they express CD16 but not CD1c (Fig. 7D). Meanwhile, CD1c^low-int^ CD14^low^ cells are heterogenous and T cells were a major population of them. Notably, quantitative real-time PCR analysis among fractions of CD45^+^ PECs revealed that expression of *GATA6* was found only in CD1c^low-int^ CD14^low^ cells (Fig. 7B), indicating that *GATA6*-expressing macrophages are contained in CD1c^low-int^ CD14^low^ cells. According to the scRNA-seq analysis, it can be assumed that CD9 expression can be used to identify *GATA6*-expressing macrophages. Then, we have examined surface CD9 expression in CD1c^low-int^ CD14^low^ cells in CAPD or LC patient-derived PECs. The frequency of CD1c^low-int^ CD14^low^ CD9^+^ cells was decreased in LC patient-derived PECs (Fig.7C).

**FIG. 7.**
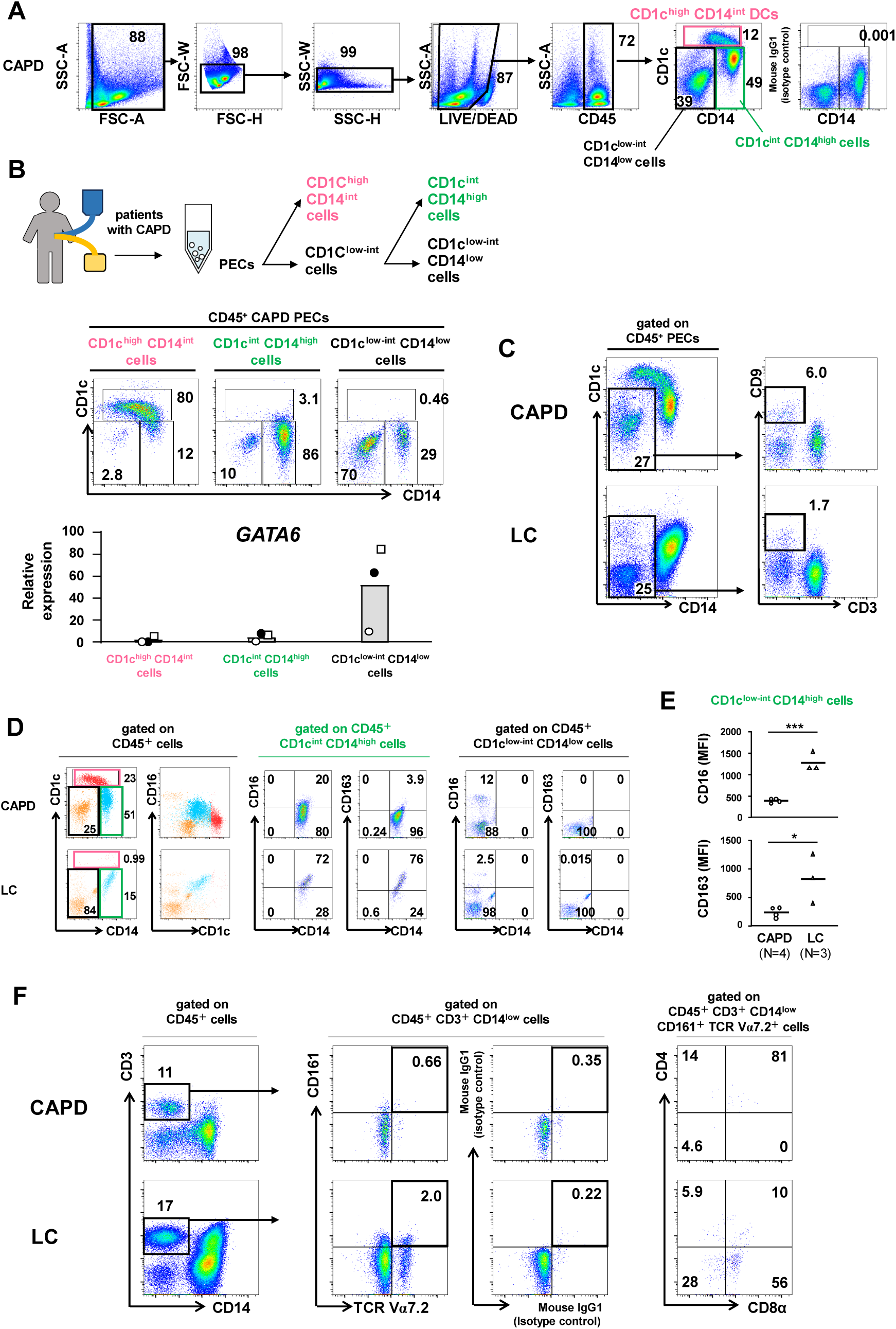
Flowcytometry analysis revealed decrease of DCs and increase of CD16^high^ CD163^high^ macrophages and CD8α^+^ MAIT cells in LC PECs. (A) Gating strategy to identify CD1c^high^ CD14^int^, CD1c^int^ CD14^high^ and CD1c^low-int^ CD14^low^ cells in CAPD PECs. Numbers indicate percentages of gated cells. (B) CD1c^high^ CD14^int^, CD1c^int^ CD14^high^ and CD1c^low-int^ CD14^low^ cells were purified from CAPD PECs by MACS system. Expression of *GATA6* among these three subsets is shown (N=3). (C) Frequency of CD1c^low-int^ CD14^low^ CD9^+^ cells in CAPD PECs and LC PECs. Numbers indicate percentages of gated cells. (D) Surface expression of CD1c, CD14, CD16 and CD163 in CD1c^high^ CD14^int^, CD1c^int^ CD14^high^ and CD1c^low-int^ CD14^low^ cells in CAPD PECs or LC PECs. Red dots indicate CD1c^high^ CD14^int^ cells, blue dots indicate CD1c^int^ CD14^high^ cells and orange dots indicate CD1c^low-int^ CD14^low^ cells, respectively. Numbers indicate percentages of gated cells. (E) Surface expression of CD16 and CD163 in CD1c^low-int^ CD14^high^ cells in CAPD PECs (N=4) or LC PECs (N=3). MFI indicates mean fluorescence intensity. Statistical significance was determined by an unpaired two-tailed Student’s t test. *p* values are indicated by \**p* < 0.05, \*\*\**p* < 0.001. *p* < 0.05 was considered statistically significant. (F) Surface expression of T cell markers in CAPD PECs or LC PECs. Numbers indicate percentage of gated cells.

Then we have examined the fluctuation of immune cell populations in LC patient-derived PECs. CD1c^high^ CD14^int^ cells, that is, DCs were present in CAPD patient-derived PECs, while they were decreased or absent in LC patient-derived PECs (Fig.7D). CD1c^int^ CD14^high^ cells were decreased in LC patient-derived PECs compared with in CAPD patient-derived PECs. We have then analyzed expression of CD16 and CD163 in CD1c^int^ CD14^high^ cells (Fig. 7E). Expression of CD16 as well as CD163 was higher in LC patient-derived PECs than in CAPD patient-derived PECs. Meanwhile, expression of CD16 and CD163 on DCs and CD1c^low-int^ CD14^low^ cells remained low in LC patient-derived PECs. Thus, expression of CD16 and CD163 was elevated only in CD1c^int^ CD14^high^ cells among LC patient-derived PECs. In patients with LC CP-B, T cells including CD161^+^ TCR Vα7.2^+^ MAIT cells were increased (Fig.7F). These results were consistent with both scRNA-seq analysis and bulk RNA-seq analysis.

## DISCUSSION

The current study is the first report on comparative analysis to characterize immune cell profiles of PECs between patients with LC and patients undergoing CAPD. Five main cell clusters, including T cells, B cells, DCs and *GATA6*- and *FCGR3A*-expressing macrophages, were found in PECs. In CAPD patient-derived PECs, DCs and *GATA6*- and *FCGR3A* expressing macrophages were dominant cell populations. This immune cell profile is consistent with previous reports.^11,12^ Meanwhile, in LC patient-derived PECs, DCs and two clusters of macrophages decreased and T cells became dominant. Among DCs and macrophages, *FCGR3A*-expressing macrophages were dominant in LC patient-derived PECs. Bulk RNA-seq analysis further clarified up- or down-regulated gene lists between PECs from patients with LC and those from patients undergoing CAPD. The results were consistent with scRNA-seq analysis of PECs.

*FCGR3A*-expressing macrophages express *CD163*, *MARCO* and *LYVE1* and represent M2 type and anti-inflammatory phenotype. Meanwhile, DCs, which express *CD1C* and *CD1E*, show M1 type and proinflammatory phenotype. Therefore, LC PECs tend to be skewed to M2. In the liver, MAIT cells can promote liver fibrosis through the recruitment of profibrotic monocytes.^14^ In this context, increase of MAIT cells in the peritoneal cavity might cause recruitment of *FCGR3A*-expressing macrophages. It should be intriguing to clarify the roles of such recruited *FCGR3A*-expressing macrophages in progression of liver fibrosis.

*GATA6*-expressing macrophages are resident in the peritoneal cavity and also profibrotic, as they express collagen gene such as *COL1A1* and repair-related genes (*THBS1*, *TGFB2*) (Fig.2A-F). *GATA6*-expressing macrophages represent a minor population in LC patient-derived PECs. However, it is still possible that such macrophages migrate to the liver and provoke fibrosis, because *GATA6*-expressing macrophages in the peritoneal cavity are required for liver protection in murine liver injury model.^3^

We have further characterized gene expressions with LC severity. Compared with PECs from patients undergoing CAPD, expression of six genes was decreased in PECs from patients with LC CP-B and their expression levels were further reduced in PECs from patients with LC CP-C. Those genes were expressed mainly in DCs and *GATA6*-expressing macrophages. Among those genes, *CTLA4* encodes CTLA4, which can inhibit T cell proliferation.^15^ *C15orf48* encodes modulator of cytochrome C oxidase during inflammation, which is a mitochondrial protein critically involved in autophagy induction and *C15orf48*-deficient mice showed development of autoimmunity, indicating that *C15orf48* regulates immune functions.^16^ Decreased expression of these immunosuppressive molecules might perturb peritoneal immune homeostasis in patients with LC.

Meanwhile, four genes (*IDH1*, *CMKLR1*, *SLC7A8* and *SLC46A*) were found to increase in their expression as the LC stage progresses. Those genes were highly expressed in *GATA6*- and *FCGR3A*-expressing macrophages. *IDH1* encodes IDH, which generates α-ketoglutarate from isocitrate, thereby promoting differentiation of macrophages towards M2.^17,18^ Expression of *CMKLR1* is high in macrophages from broncho alveolar lavages of patients with cystic fibrosis, indicating involvement of *CMKLR1* in fibrosis.^19^ Expression of *SLC7A8* is also correlated with that of CD163 in M2 macrophages.^20^ Thus, elevated expression of these genes might be related to upregulation of fibrotic or M2 type macrophage functions. It should be interesting whether or how those gene products contribute to progression of liver fibrosis.

Expression of some genes was elevated in PECs from patients with LC CP-B, but decreased in PECs from patients with LC CP-C. KEGG pathway analysis of such gene enrichment of the ‘Natural killer cell mediated cytotoxicity’ pathway was enriched (Fig. 6C). This enrichment can be assumed to be due to increase of T cells (Supplementary Figure 3A,B). Especially cytotoxic T cell signature genes, *GZMB* and *IFNG,* were prominently elevated. MAIT cells, which are also cytotoxic, are decreased in peripheral bloods, but increased in PECs from patients with decompensated LC and spontaneous bacterial peritonitis (SBP).^2,21^ In that study, patients with LC were not categorized by CP classification. Meanwhile, we studied patients with LC at CP-B and CP-C in the absence of SBP (Supplementary Table S1). It is interesting to elucidate how T cells including MAIT cells are increased at CP-B and then decreased at CP-C. CD163^high^ M2 type macrophages can inhibit T cell proliferation.^22^ Therefore, the T cell decrease can be ascribed to dominancy of M2 macrophages at CP-C. It is also possible that activated T cells leave the peritoneal cavity and migrate to the liver.

FACS analysis showed that, in LC patient-derived PECs, CD1c^high^ CD14^int^ DCs were decreased, while CD16^high^ CD163^high^ cells were increased (Fig. 7D,E). We further found that CD1^low-int^ CD14^low^ cells include *GATA6*-expressing macrophages (Fig. 7B) and that these cells were decreased in LC patient-derived PECs. Furthermore, T cells including MAIT cells were increased and existed as a major population in LC patient-derived PECs. These flowcytometry analysis results were consistent with the results of scRNA-seq and bulk RNA-seq analyses.

We thus characterized immune cell profiles of PECs from patients with LC. Whereas DCs and macrophages were dominant in PECs from patients undergoing CAPD, T cells were the main population in PECs from patients with LC. Although DCs and macrophages were decreased in PECs from patients with LC, *FCGR3A*-expressing macrophages, which are profibrotic, were relatively dominant over DCs and *GATA6*-expressing macrophages. We also compared gene expression profiles between CP-B and CP-C and clarified that T cells were increased at CP-B but decreased at CP-C. We further identified a set of genes expression of which was increased or decreased with LC severity. Our present findings should have a potential to lead to further development of diagnosis or therapeutic maneuver for LC.

## ACKNOWLEDGEMENTS

We thank Ms. Sumiko Okura and Ms. Aoi Matsumura-Tawaki for secretarial assistance. We acknowledge proofreading and editing by Benjamin Phillis at the Clinical Study Support Center, Wakayama Medical University.

---

**Supplementary Figure 1.**
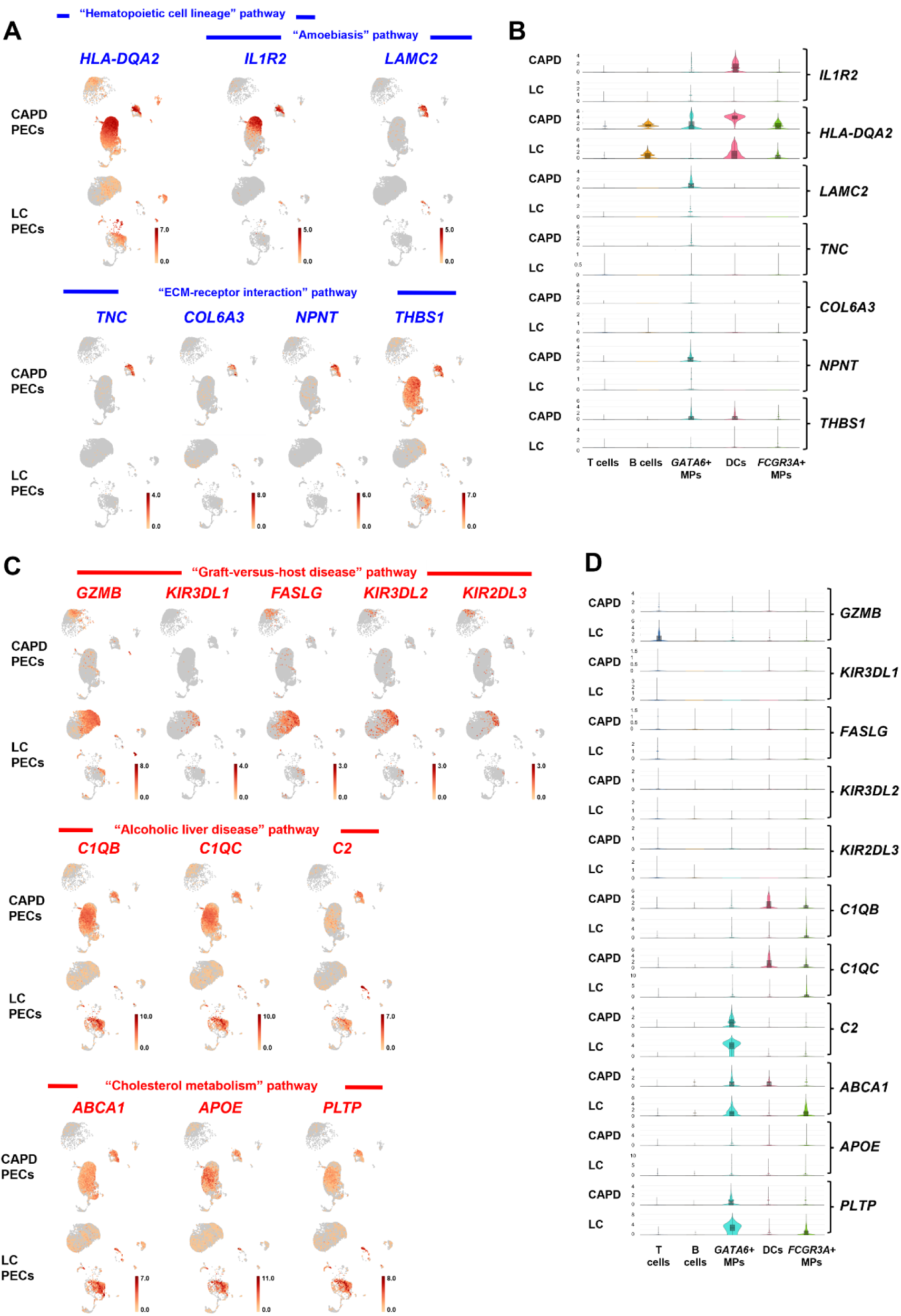
Gene expression profiles of up- or down-regulated genes in LC patient-derived PECs by scRNA-seq analysis. (A,B) Expression of *HLA-DQA2*, *IL1R2*, *LAMC2*, *TNC*, *COL6A3*, *NPNT* and *THBS1* is shown in feature plots (A) and violin plots (B). (C,D) Expression of *GZMB*, *KIR3DL1*, *FASLG*, *KIR3DL2*, *KIR2DL3*, *C1QB*, *C1QC*, *C2*, *ABCA1*, *APOE* and *PLTP* is shown in feature plots (C) and violin plots (D).

**Supplementary Figure 2.**
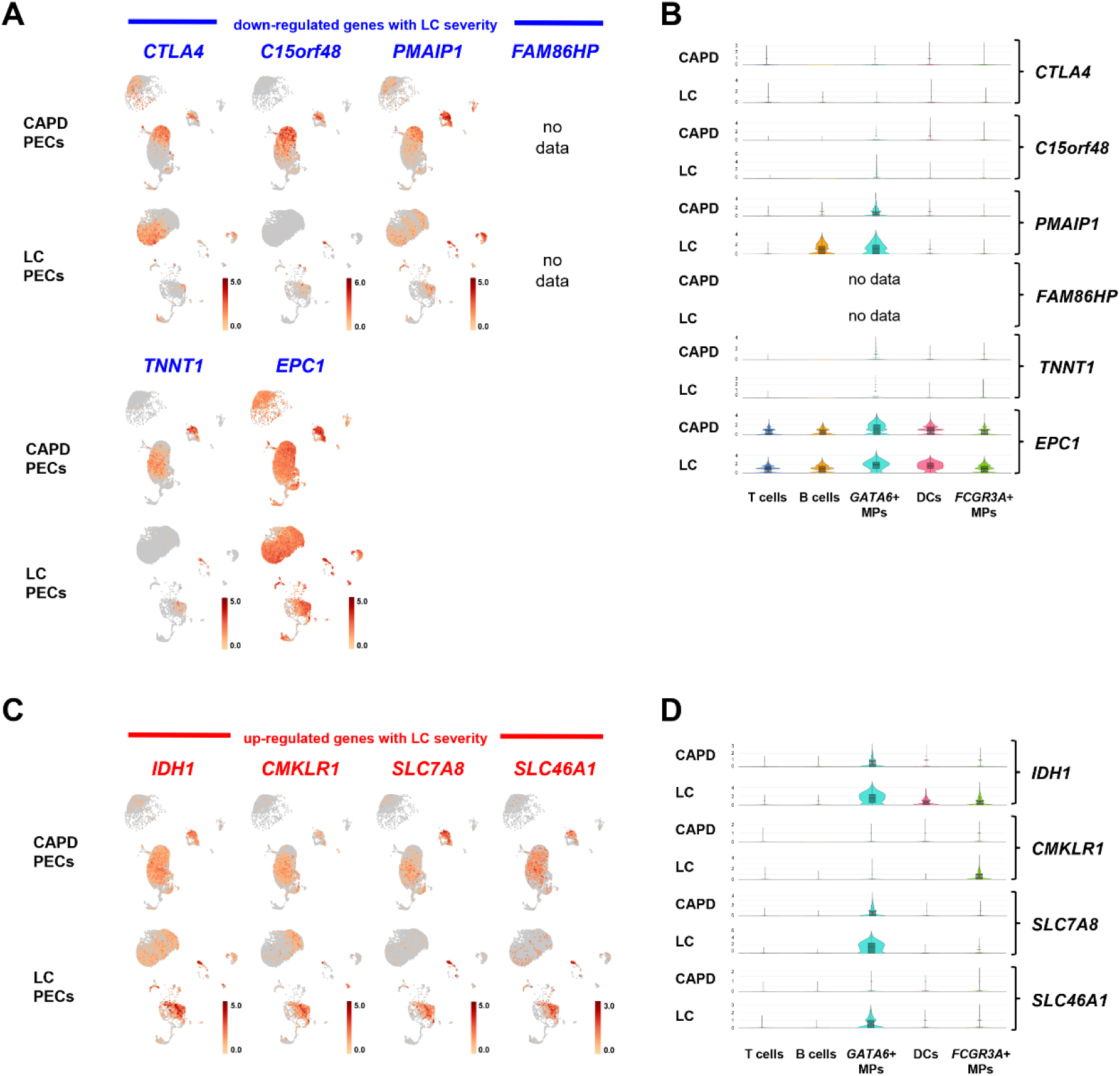
Gene expression profiles of up- or down-regulated genes with LC severity by scRNA-seq analysis. (A,B) Expression of *CTLA4*, *C15orf48*, *PMAIP1*, *FAM86HP*, *TNNT1* and *EPC1* is shown in feature plots (A) and violin plots (B). (C,D) Expression of *IDH1*, *CMKLR1*, *SLC7A8* and *SLC46A1* is shown in feature plots (C) and violin plots (D).

**Supplementary Figure 3.**
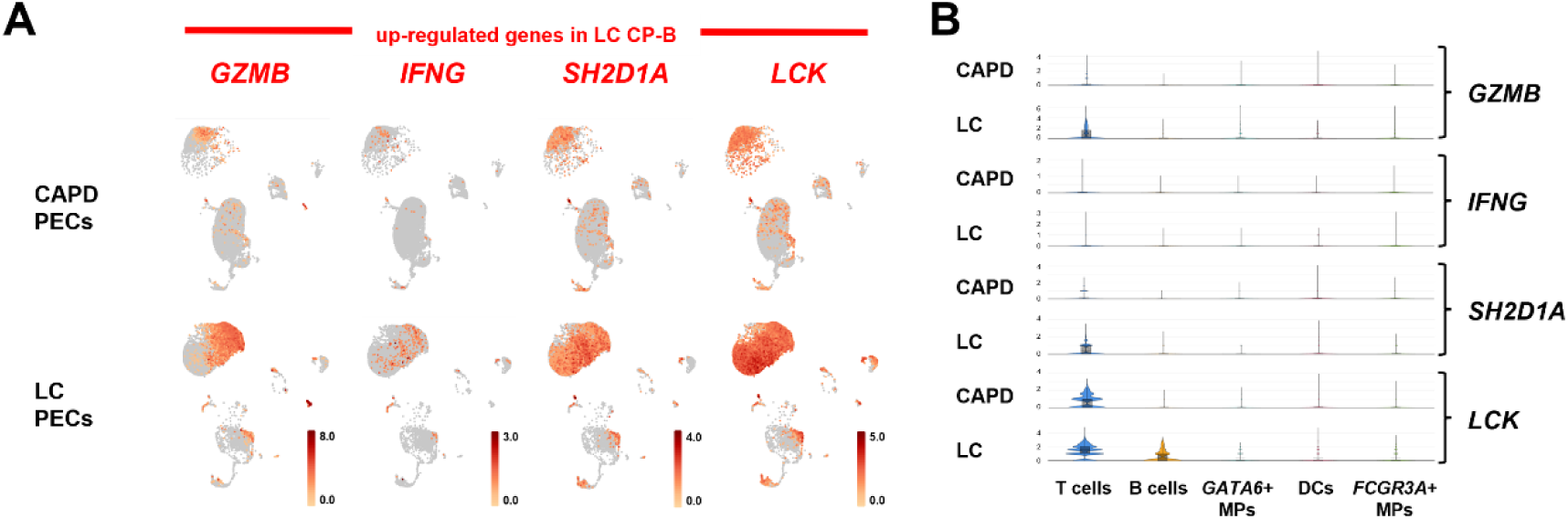
Gene expression profiles of upregulated genes in LC CP-B patient-derived PECs by scRNA-seq analysis. (A,B) Expression of *GZMB*, *IFNG*, *SH2D1A* and *LCK* is shown in feature plots (A) and violin plots (B).

**Supplementary Table S1.**
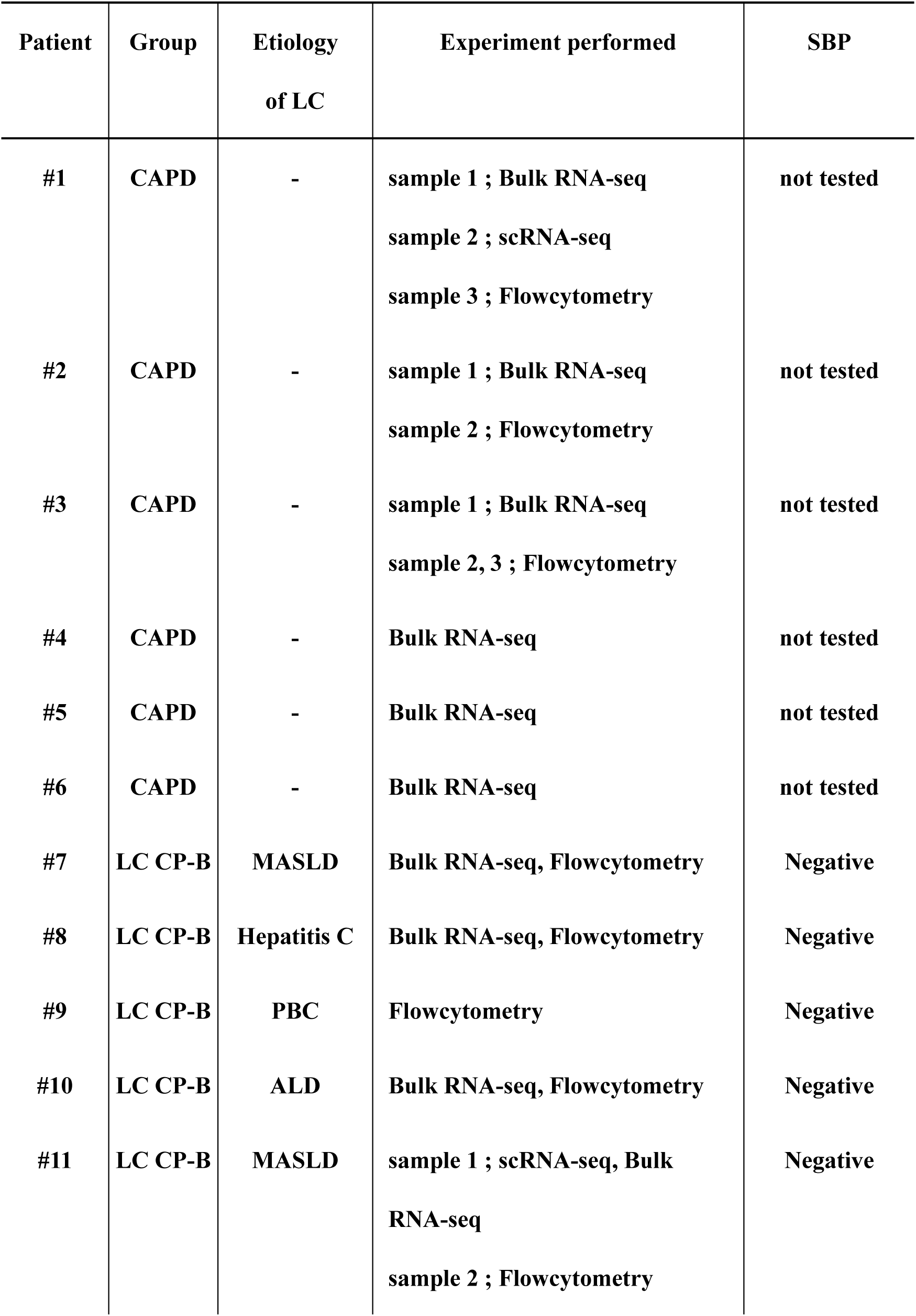
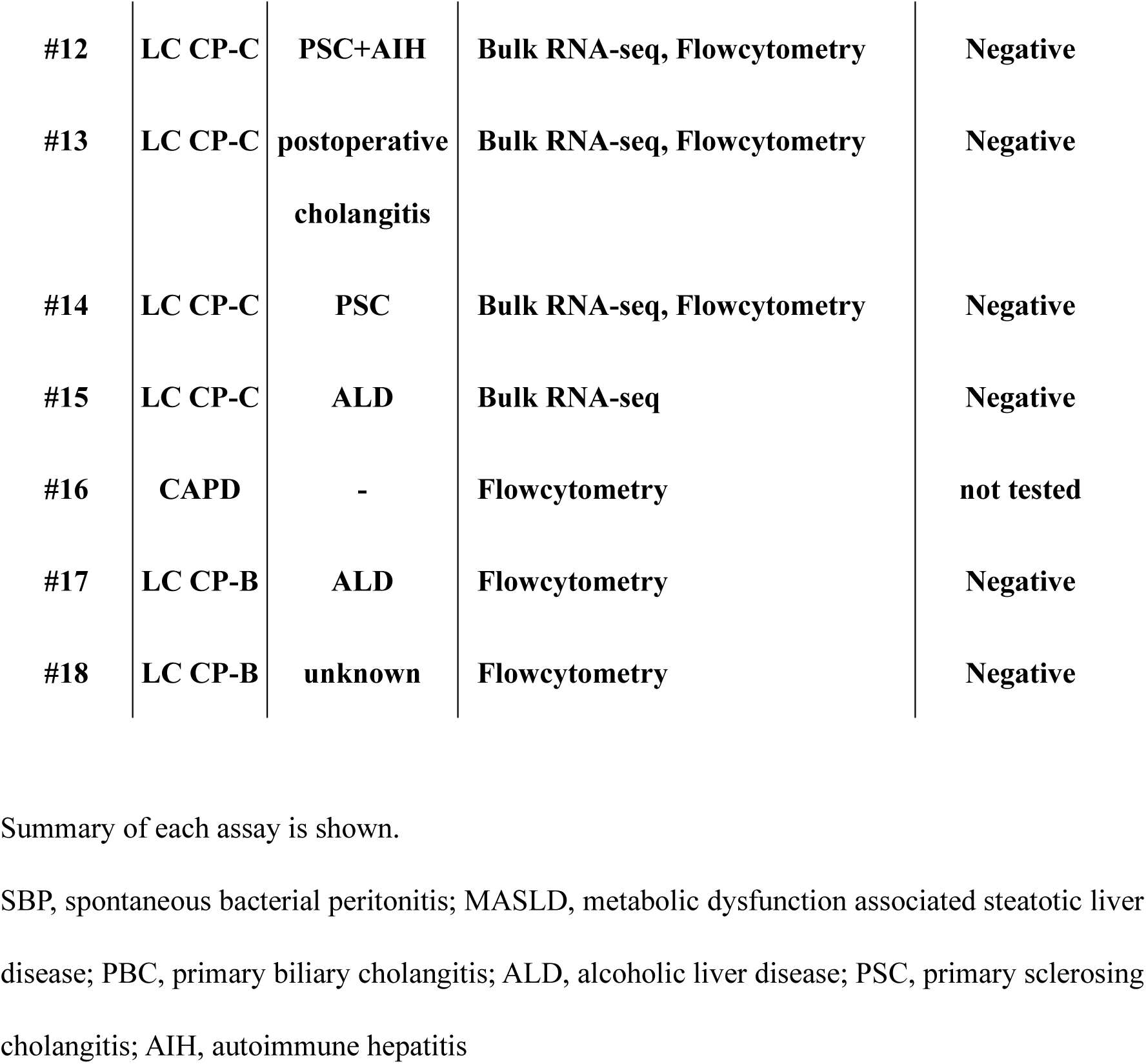
Patients undergoing CAPD or with LC enrolled in this study.

| **Patient** | **Group** | **Etiology**<br>**of LC** | **Experiment performed** | **SBP** |
| --- | --- | --- | --- | --- |
| **#1** | **CAPD** | **-** | **sample 1 ; Bulk RNA-seq**<br>**sample 2 ; scRNA-seq**<br>**sample 3 ; Flowcytometry** | **not tested** |
| **#2** | **CAPD** | **-** | **sample 1 ; Bulk RNA-seq**<br>**sample 2 ; Flowcytometry** | **not tested** |
| **#3** | **CAPD** | **-** | **sample 1 ; Bulk RNA-seq**<br>**sample 2, 3 ; Flowcytometry** | **not tested** |
| **#4** | **CAPD** | **-** | **Bulk RNA-seq** | **not tested** |
| **#5** | **CAPD** | **-** | **Bulk RNA-seq** | **not tested** |
| **#6** | **CAPD** | **-** | **Bulk RNA-seq** | **not tested** |
| **#7** | **LC CP-B** | **MASLD** | **Bulk RNA-seq, Flowcytometry** | **Negative** |
| **#8** | **LC CP-B** | **Hepatitis C** | **Bulk RNA-seq, Flowcytometry** | **Negative** |
| **#9** | **LC CP-B** | **PBC** | **Flowcytometry** | **Negative** |
| **#10** | **LC CP-B** | **ALD** | **Bulk RNA-seq, Flowcytometry** | **Negative** |
| **#11** | **LC CP-B** | **MASLD** | **sample 1 ; scRNA-seq, Bulk**<br>**RNA-seq**<br>**sample 2 ; Flowcytometry** | **Negative** |
| **#12** | **LC CP-C** | **PSC+AIH** | **Bulk RNA-seq, Flowcytometry** | **Negative** |
| **#13** | **LC CP-C** | **postoperative**<br>**cholangitis** | **Bulk RNA-seq, Flowcytometry** | **Negative** |
| **#14** | **LC CP-C** | **PSC** | **Bulk RNA-seq, Flowcytometry** | **Negative** |
| **#15** | **LC CP-C** | **ALD** | **Bulk RNA-seq** | **Negative** |
| **#16** | **CAPD** | **-** | **Flowcytometry** | **not tested** |
| **#17** | **LC CP-B** | **ALD** | **Flowcytometry** | **Negative** |
| **#18** | **LC CP-B** | **unknown** | **Flowcytometry** | **Negative** |

Summary of each assay is shown.
SBP, spontaneous bacterial peritonitis; MASLD, metabolic dysfunction associated steatotic liver
disease; PBC, primary biliary cholangitis; ALD, alcoholic liver disease; PSC, primary sclerosing
cholangitis; AIH, autoimmune hepatitis

**Supplementary Table S2.** Clinical characteristics of the study individuals for scRNA-seq analysis.

| Analysis of scRNA-seq |  |  |
| --- | --- | --- |
| Characteristic | CAPD (N = 1) | LC CP-B (N = 1) |
| Age, y | 69 | 76 |
| Male sex | Male | Male |
| Serum leukocytes ($\times 10^9/l$) | 5.86 | 3.09 |
| Plt ($\times 10^9 /l$) | 248 | 52 |
| Serum PT (%) | - | 60.9 |
| Serum Albumin (g/dl) | 3.0 | 2.9 |
| AST (U/l) | 13 | 15 |
| ALT (U/l) | 13 | 11 |
| T-Bil (mg/dl) | 0.5 | 0.9 |

**Supplementary Table S3.**
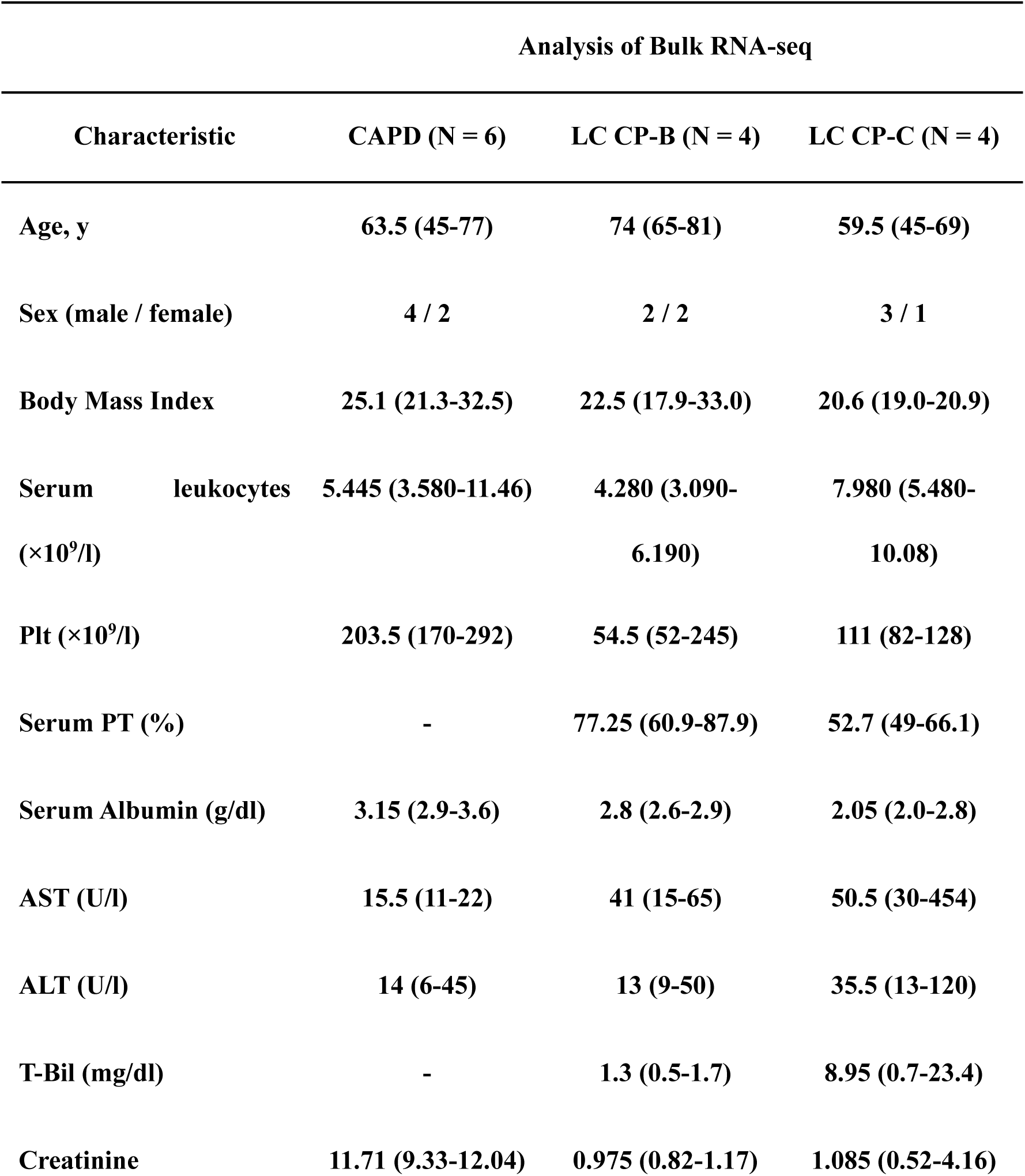
Clinical characteristics of the study individuals for bulk RNA-seq analysis.

**Supplementary Table S4.** Antibodies used for flowcytometry analysis.

| Antibody name | Color | Clone | Cat | Antibody<br>source |
| --- | --- | --- | --- | --- |
| anti-CD45 | APC | HI30 | 304037 | BioLegend |
| anti-CD1c | Biotin | L161 | 331504 | BioLegend |
| anti-CD14 | PE | 61D3 | 12-0149-42 | Invitrogen |
| anti-CD9 | PerCPCy5.5 | HI9A | 312110 | BioLegend |
| anti-CD3 | APCCy7 | UCTH1 | 300426 | BioLegend |
| anti-CD16<br>(encoded by *FCGR3A*) | Pacific Blue | 3G8 | 980106 | BioLegend |
| anti-CD163 | FITC | GHI/61 | 333617 | BioLegend |
| anti- CD161 | PECy7 | HP-3G10 | 339918 | BioLegend |
| anti-TCR Vα7.2 | Brilliant violet 421 | 3C10 | 351716 | BioLegend |
| anti-CD4 | PerCPCy5.5 | RPA-T4 | 300530 | BioLegend |
| anti-CD8α | FITC | RPA-T8 | 301060 | BioLegend |

